# Genomes of the extinct Bachman’s Warbler shows high divergence and no evidence of admixture with other extant *Vermivora* Warblers

**DOI:** 10.1101/2022.12.20.521272

**Authors:** Andrew W. Wood, Zachary A. Szpiech, Irby Lovette, Brian Tilson Smith, David P. L. Toews

**Affiliations:** Department of Biology, 619 Mueller Laboratory, Pennsylvania State University, University Park, Pennsylvania 16802, USA; Institute for Computational and Data Sciences, Pennsylvania State University, University Park, Pennsylvania 16802, USA; Fuller Evolutionary Biology Program, Cornell Lab of Ornithology, Cornell University, 159 Sapsucker Woods Road, Ithaca, NY 14850, USA; Department of Ornithology, American Museum of Natural History, Central Park West at 79th Street, New York, NY 10024, USA

## Abstract

Bachman’s Warbler (*Vermivora bachmanii*) – last sighted in 1988 – is one of the few North American passerines that have gone extinct. Given the extensive ongoing hybridization of its two extant congeners – the Blue-Winged Warbler (*V. cyanoptera*) and Golden-Winged Warbler (*V. chrysoptera*) – and shared patterns of plumage variation between Bachman’s Warbler and hybrids between those extant species, it has been suggested that Bachman’s Warbler might have also had a component of hybrid ancestry. Here, we use historic DNA (hDNA) and whole genome sequencing of Bachman’s Warblers collected at the turn of the 20^th^ century to address this possibility. We combine these data with genomes of the two extant *Vermivora* species to examine patterns of population differentiation, inbreeding, and gene flow. In contrast to the admixture hypothesis, the genomic evidence is consistent with *V. bachmanii* being a highly divergent, reproductively isolated species, with no evidence of introgression. We show that both *V. bachmanii* and *V. chrysoptera* have elevated runs of homozygosity compared to *V. cyanoptera*, consistent with the effects of a small effective population size or population bottlenecks in the former two species. We also found—using population branch statistic estimates of all three species—previously undocumented evidence of lineage-specific evolution in *V. chrysoptera* near a novel pigmentation gene candidate for warblers, *CORIN*, which is a known modifier of *ASIP*, which is in turn involved in melanic throat and mask coloration in this family of birds. Together, these genomic results also highlight how natural history collections are such invaluable repositories of information about extant and extinct species.

**Significance:** Few common North American passerines have gone extinct. Bachman’s Warbler is, unfortunately, one that has—the last sighting was in 1988. Here we use whole genome historical DNA from museum specimens of Bachman’s warblers collected at the turn of the 20^th^ century to learn about the evolution of this species and test whether there was evidence for hybridization and gene flow between it and two extant members of the same genus which, today, hybridize extensively. We find Bachman’s warbler was highly divergent with no evidence of gene flow. We also find evidence of elevated “runs of homozygosity” in both Bachman’s warbler and one of the two extant *Vermivora* species, suggesting the effects of a small population size or population bottlenecks.

## Introduction

*“Within an area of ten or fifteen acres there must have been nearly one thousand warblers, of which probably five per cent were Bachman’s. It was comparatively easy to identify them   and with more time I could have taken thrice as many specimens as were actually obtained.”* (Brewster 1891)

Few locally common North American passerines have gone extinct. Bachman’s Warbler (*Vermivora bachmanii*) is, unfortunately, one of the few that almost certainly has, as the last confirmed sighting of this songbird was in 1988 in Louisiana (Hamel 2020). Subsequently, many targeted search efforts have failed to locate any individuals, and it has not been observed despite the multifold increase in birding activity over recent decades. It is therefore widely accepted to be extinct and was officially declared so in 2021 by the United States Fish and Wildlife Service (Department of the Interior 2021).

The only extant members of the same genus, *Vermivora*, are the Golden-Winged Warbler (*Vermivora chrysoptera*) and the Blue-Winged Warbler (*Vermivora cyanoptera*), which are notable among parulid warblers for their widespread and extensive hybridization (Gill 1980, Shapiro et al. 2004). This includes the “named” hybrid morphs “Lawrence’s” and “Brewster’s” Warblers, which were originally each thought to be a distinct species before they were found to be hybrids between *V. cyanoptera* and *V. chrysoptera* (Faxon 1913). Genomic analyses of *V. cyanoptera* and *V. chrysoptera* (and their hybrids) have since revealed few and relatively small regions of the genomes that are divergent between them. These small divergent regions primarily house pigmentation genes, and they presumably underlie the markedly different plumage phenotypes of these species (Toews et al. 2016, Baiz et al. 2020). For instance, the distinctive black throat patch and face mask of *V. chrysoptera* appears in the “Lawrence’s” Warbler hybrid phenotype but is absent in the “Brewster’s” hybrid type (Fig 1A), and this difference is associated with a 10kb chromosomal region upstream of the *Agouti signaling protein (ASIP*) gene, which has a gene product involved in the melanogenesis pathway (Toews et al. 2016, Baiz et al. 2021). This black throat patch is also one of the most distinctive plumage traits of Bachman’s Warbler, making it phenotypically parallel to *V. chrysoptera* and the "Lawrence’s" hybrid form (at least for this one trait).

**Figure 1.**
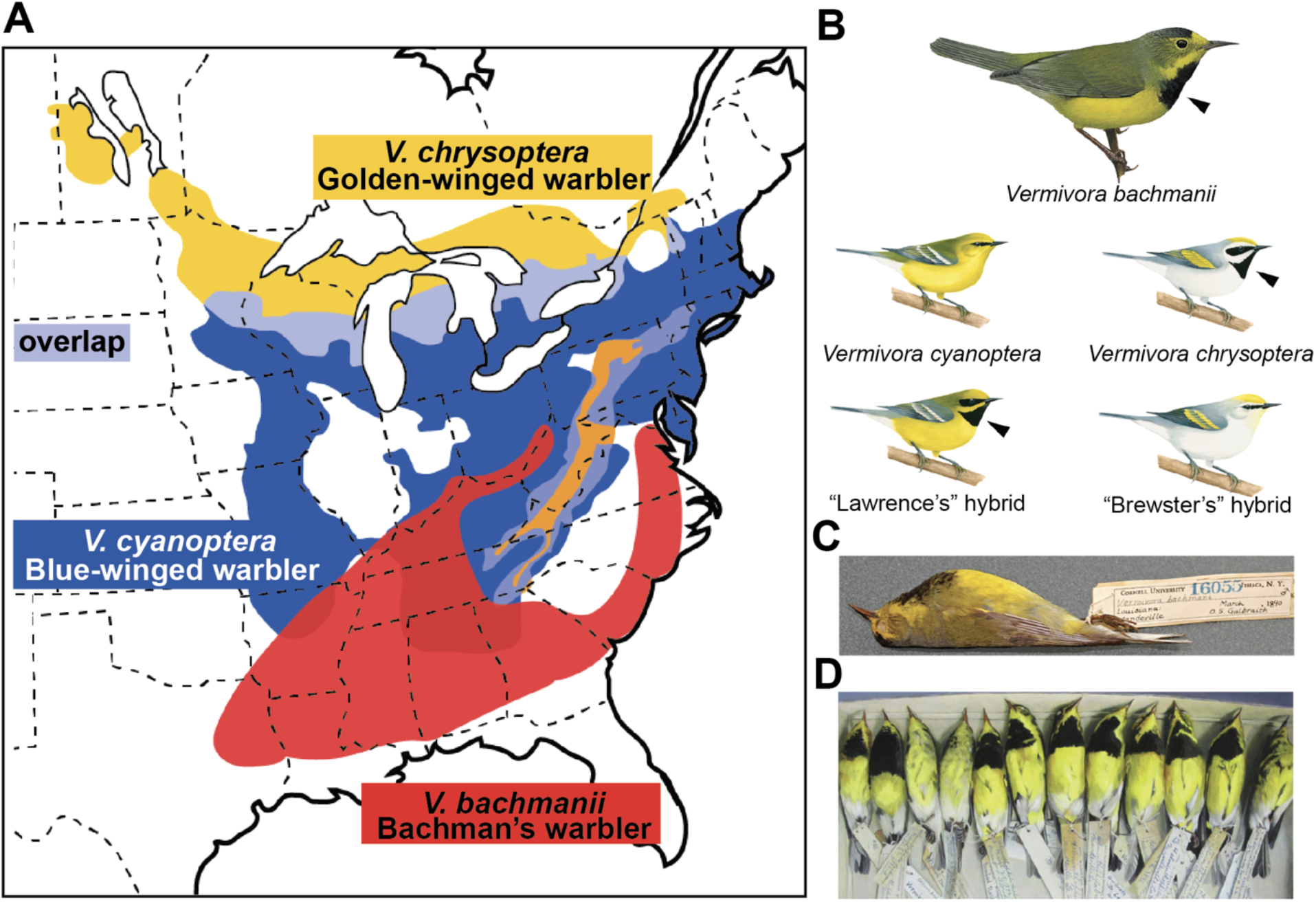
Map (A) and plumage phenotypes (B-D) of *Vermivora* Warblers. The range of *V. bachmanii* inferred from Hamel (2020). Bachman’s warbler illustration used with permission © Lynx Edicions. Other *Vermivora* illustrations by Liz Clayton Fuller. Photo in (C) from Cornell University Museum of Vertebrates, (D) is an oil painting by Isabella Kirkland (2011) with references from specimens at the AMNH.

The current extensive hybridization between the two extant *Vermivora* species, combined with the similarity of plumage patches between *V. bachmanii* and contemporary *Vermivora* hybrids, caused Hamel (2018) to ask whether *V. bachmanii* might have arisen via hybridization. Given the known distribution and habitat preference of *V. bachmanii* (bamboo, Remsen 1986), it seems unlikely that it co-occurred in breeding sympatry with *V. chrysoptera*, and even perhaps not in tight local sympatry with *V*. *cyanoptera* (Figure 1A). However, extant *Vermivora* live in early successional, secondary growth habitat, which is both geographically and temporally dynamic on the landscape. Thus, it may have presented an opportunity for local sympatry for two or more of the species and possibly resulted in hybridization.

Genetic studies of the question of possible hybridization and introgression in *V. bachmanii* have not previously been possible. Prior to 2021, a small fragment of the mitochondrial gene ND2 obtained by Lovette et al. (2010) was the only DNA sequence available for *V. bachmanii*, and the maternal inheritance of mtDNA makes this genetic marker inadequate to detect genomic introgression. Likewise, a sequence-capture dataset from ultra-conserved element loci produced insights into the demographic history of *V. bachmanii* (Smith et al. 2021) but is unsuitable to study past hybridization among extant *Vermivora* because their genomes are so uniformly similar (Toews et al. 2016).

Here we generate the data necessary to study the possibility of historical admixture among *Vermivora*. We use whole genome data for *V. bachmanii* produced from the toepad DNA of seven specimens collected in the southern USA between 1889 and 1915 (Fig 1B). We compared patterns of divergence among the three species to quantify the extent and distribution of divergence across *Vermivora* Warblers, as well as to test for signals of introgression. We also explore genomic patterns relevant to the decline and conservation of small populations—runs of homozygosity (ROH)—among all three *Vermivora* species; ROH are long stretches of identical-by-descent haplotypes within individual genomes and are indicators of small population size, bottlenecks, and inbreeding (Ceballos et al. 2018; Kardos et al. 2015).

## Methods

We generated genomic data from *V. bachmanii* from toepads sampled from museum skins from the Cornell Museum of Vertebrates (CUMV; *n* = 2) and the American Museum of Natural History (AMNH; *n* = 6). Six sampled birds were male and two were female (Table 1). Toepad specimens were extracted in dedicated labs for working on historical samples at the AMNH and CUMV. For the pilot study (CUMV samples), we used a phenol-chloroform DNA extraction method. For the AMNH samples, we used the Qiagen DNeasy Blood and Tissue Kit with several modifications to improve DNA yields that included washing each toe pad sample with H2O and EtOH, a longer digestion time, and using QIAquick PCR filter columns to size select for smaller fragments of DNA. One of the two samples from the CUMV did not result in any useable genomic data and was therefore excluded from downstream analyses.

**Table 1.** Sample information for the seven hDNA *Vermivora bachmanii* samples.

| Species | Sample ID | Museum | Sex | Year | Location | Coverage (X) |
| --- | --- | --- | --- | --- | --- | --- |
| *V. bachmanii* | 229845 | AMNH | male | 1907 | Missouri | 4.4 |
| *V. bachmanii* | 49494 | AMNH | male | 1890 | Florida | 4.4 |
| *V. bachmanii* | 759210 | AMNH | male | 1891 | Louisiana | 4.2 |
| *V. bachmanii* | 759214 | AMNH | male | 1915 | South Carolina | 4.6 |
| *V. bachmanii* | 759216 | AMNH | female | 1889 | Florida | 3.4 |
| *V. bachmanii* | 759220 | AMNH | female | 1890 | Florida | 4.7 |
| *V. bachmanii* | 16055 | CUMV | male | 1890 | Louisiana | 0.795 |

The samples were sequenced across two separate NextSeq lanes. The CUMV samples were sequenced in 2018 as part of a pilot project conducted at the Cornell Institute for Biotechnology. The AMNH samples were sequenced in 2022 at the Penn State Genomics Core facility. We followed the protocol for the Illumina TruSeq Nano library preparation kit, with two important modifications. First, given the DNA was already degraded, we did not mechanically shear the DNA. Second, we did not perform a SPRI bead size selection, with the justification that omitting size selection would result in smaller fragments, but would produce more resultant reads for analysis. Therefore, in place of the size selection step, we used undiluted SPRI beads. For sequencing, 24 individual birds from a variety of projects were individually indexed and pooled on an Illumnia NextSeq 500 lane using paired-end 150bp sequencing chemistry. The *V. bachmanii* were then bioinformatically de-multiplexed from the other individuals not related to the present study.

We combined our genome data from *V. bachmanii* with resequenced genomes of previously published extant *Vermivora:* ten *V. cyanoptera*, ten *V. chrysoptera*, and one each of the Lawrence’s and Brewster’s hybrid phenotype (Toews et al. 2016; Baiz et al. 2020). For the ABBA-BABA test (below) we also included 5X genomic data from an outgroup species, the ovenbird (*Seiurus aurocapilla; n* = 5), which is the sole member of its genus and the sister species to all other wood warblers (Lovette et al. 2010).

We first used the program AdapterRemoval (Schubert et al. 2016) to trim and collapse overlapping paired reads with the following options: “-collapse-trimns - minlength 20 -qualitybase 33.” We then used BowTie2 (Langmead and Salzberg 2012) to align reads to the new reference genome with the “very-sensitive-local” presets and set the option “-X” (the maximum fragment length for valid paired-end alignments) to 700 bp. We aligned all read data to the Myrtle Warbler (*Setophaga coronata coronata*) reference genome (Baiz et al. 2021). We then used PicardTools (2.20.8; https://broadinstitute.github.io/picard/) to mark PCR duplicates. We quantified genomic coverage using “qualimap” (Okonechnikov et al. 2016).

We first confirmed the identity of our *V. bachmanii* specimens by comparing them to a short fragment of mitochondrial DNA, the *ND2* gene obtained for *V. bachmanii* by Lovette et al. (2010), which showed high similarity to our samples. For some of our analyses we focus on six previously identified genomic regions of divergence between *V. cyanoptera* and *V. chrysoptera* (Toews et al. 2016). This is because, outside of these regions, *V. cyanoptera* and *V. chrysoptera* genomes are effectively homogenized (Toews et al. 2016). We used PCAngsd (version 0.981; Meisner and Albrechtsen 2018) to conduct principal component analysis on the genotype likelihood data for each region separately and plotted the first two eigenvectors. Within ANGSD, we first generated a “beagle” input file using the flags “-GL 2 -doGlf 2 -doMajorMinor 1 -SNP_pval 1e-6 - doMaf 1” and then ran PCAngsd with the default parameters.

To compare patterns of divergence among the three species across the entire genome we used both *F*_ST_ estimates and population branch statistics (PBS). To estimate *F*_ST_, we used ANGSD version 0.929 (Korneliussen et al. 2014), which accounts for genotyping uncertainty in low-coverage data, and estimated *F*_ST_ for non-overlapping 10-kb windows. For each population, we calculated the site allele frequency likelihoods using the “dosaf 1” command. We then calculated the two-population site frequency spectrum and resulting *F*_ST_ estimates using “realSFS.” To calculate lineage-specific evolution (i.e., PBS) for each of the three *Vermivora* species, we used a similar approach, calculating site allele frequency likelihoods for each species, and then calculating all the pairwise two population site frequency spectra. These were then used “realSFS” with the options “-win 10000 -step 10000” to calculate the non-overlapping sliding-window PBS statistic for each of the three species.

To calculate D-statistics for the ABBA-BABA test, we used the “-doAbbababa2” function, also in ANGSD, using the options “-blockSize 100000 -do-Counts 1.” For the analysis we used *V. cyanoptera* and *V. chrysoptera* as populations one and two (H1, H2, respectively), *V. bachmanii* samples as population three (H3), and *Seiurus aurocapilla* as the outgroup (H4) using the “-useLast 1” option. A positive D statistic in this case would indicate introgression between H2 and H3.

To call ROH we used GARLIC which requires called genotypes. We first called variant sites in *V. chrysoptera* (*n*=10), *V. cyanoptera* (*n*=10), and *V. bachmanii* (*n*=6), excluding one *V. bachmanii* sample with mean coverage <1X (as well as the two hybrids) using GATK v4.2.6.1 HaplotypeCaller (Poplin et al. 2018). As these samples have low average coverage (see below), we enforced a strict filtering scheme so that we retained only the most confidently called variants. For each species, separately, we first restricted our set of sites to only biallelic SNPs on autosomes, and we set any genotype with a read depth < 6 or a genotype quality < 20 to missing. Next, we removed any sites with a missing data rate > 0.5. Finally, we filtered sites with excess heterozygosity, as these sites are likely to be sequencing errors. However, with low sample sizes statistical tests to detect excess heterozygosity have no power, so we resort to filtering any site where >= 80% of samples have a heterozygote called (a similar strategy as in Robinson et al. 2016). For *V. chrysoptera* and *V. cyanoptera* samples this means any site with 8 or more heterozygotes, and for *V. bachmanii* this means any site with 5 or more heterozygotes. These filters result in 89,207, 179,495, and 60,124 variable sites for *V. chrysoptera, V. cyanoptera*, and *V. bachmanii*, respectively. Previous work has shown is it possible to call ROH at these SNP densities (Purfield et al. 2012; Peripolli et al. 2016). We used GARLIC v1.1.6a to call ROH (Szpiech et al. 2017) as this method implements a model-based likelihood approach that incorporates genotype quality scores and handles arbitrary amounts of missing data. Each species was called separately with the following parameters set: “--max-gap 1000000, --overlap-frac 0.05, --resample 100, and --auto-winsize”. GARLIC chose the best window size for each species, resulting in a window size of 20 SNPs, 30 SNPs, and 20 SNPs for *V. chrysoptera, V. cyanoptera*, and *V. bachmanii*, respectively.

## RESULTS

Previously published *V. cyanoptera* and *V. chrysoptera* samples were sequenced to 4-5X coverage (Toews et al. 2016). The *V. bachmanii* coverage for the present study was comparable for six samples (AMNH average coverage = 4.3X), with the pilot sample having lower coverage (CUMV coverage = 0.8X; Table 1).

Patterns of genome-wide clustering demonstrate that *V. bachmanii* has had a long period of independent evolution (Figure 2). Across each of the six regions that are divergent between the other two *Vermivora* species, *V. bachmanii* consistently clusters independently along PC1 from *V. cyanoptera* and *V. chrysoptera*, which each separate along PC2. Moreover, we did not observe evidence of genomic intermediacy at any of the regions, such as manifests in the contemporary hybrid phenotypes, implying little or no historical admixture between *V. bachmanii* and either of the other two species. The non-significant result of the ABBA-BABA test is similarly consistent with this lack of admixture (across 12,201 blocks of 100kb: D = -0.0004, *P* = 0.0624).

**Figure 2.**
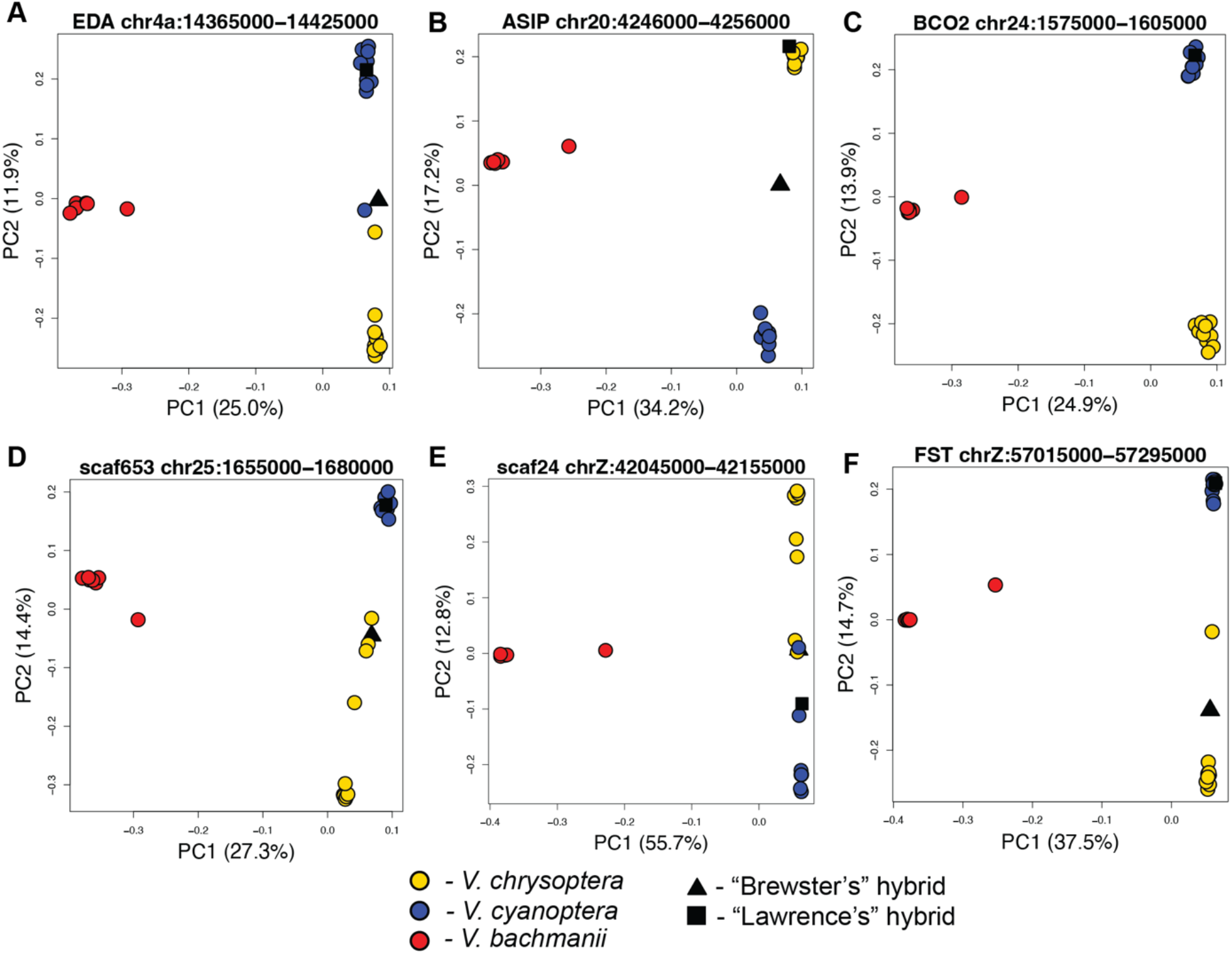
Principal components analysis of six regions of divergence identified between *V. cyanoptera* and *V. chrysoptera* using genotype likelihoods. Chromosomal coordinates of regions are shown in plot titles. Candidate pigmentation genes as in Toews et al. (2016) and, for regions without candidates, previous scaffold names are shown. (A) *EDA (ectodysplasin*) region (B) *ASIP (agouti signaling protein*) region, (C) *BCO2 (beta-carotene oxygenase 2*) region, (D) unnamed region on chromosome 25, (E), unnamed region on chromosome Z (F) *F*_ST_ (*follistatin*) region.

Patterns of divergence among the three species differed dramatically. Similar to previous studies comparing *V. cyanoptera* and *V. chrysoptera*, genome divergence between them as measured by *F*_ST_ was extremely low, save for the six small divergent regions previously identified that house pigmentation genes (Figure 3C). By contrast, each species’ pairwise comparison with *V. bachmanii* showed very high genome-wide divergence (Figures 3B and 3C).

**Figure 3.**
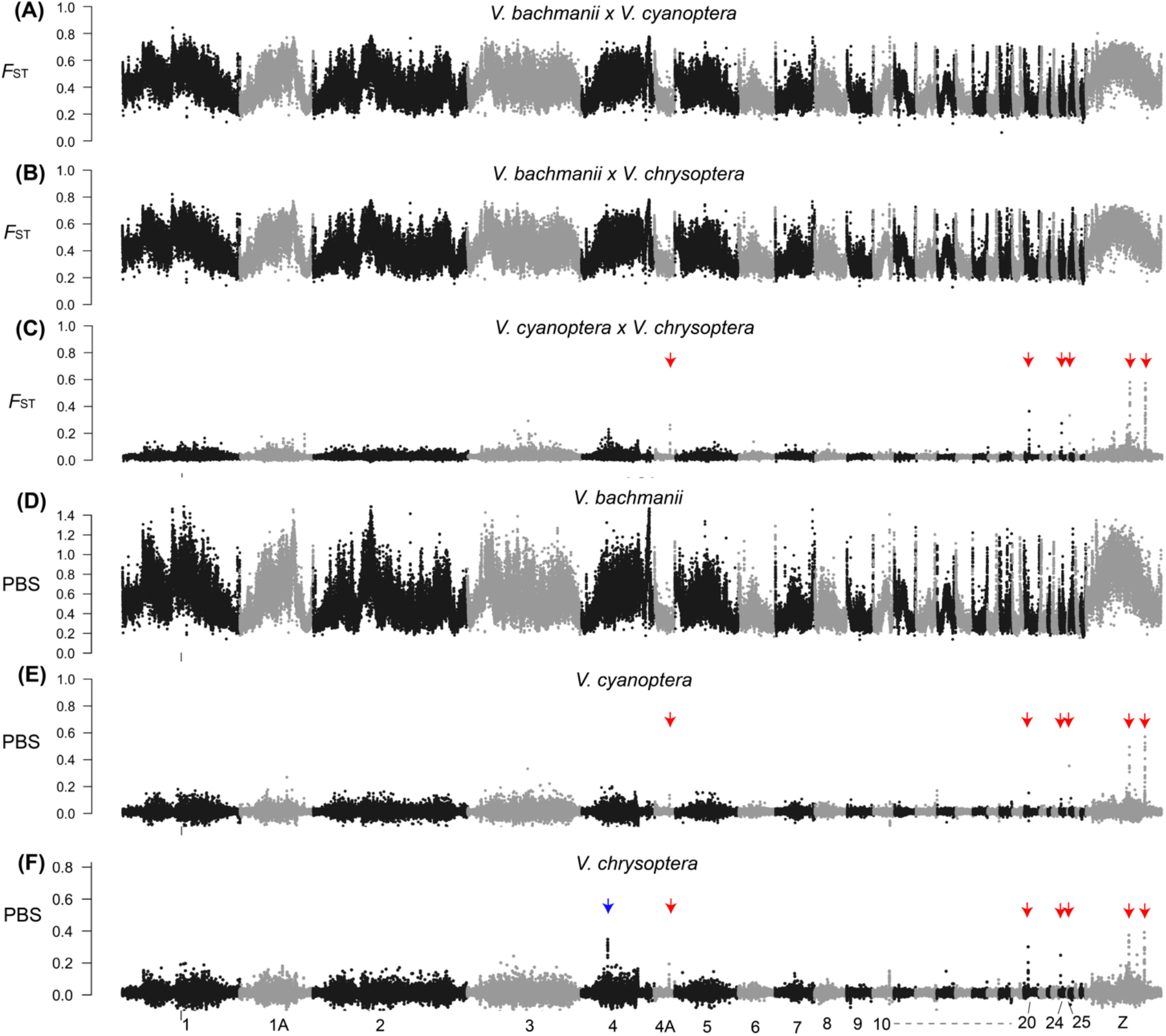
Genome-wide *F*_ST_ and PBS measures between each of the three species (A– C) as well as lineage-specific PBS estimates (D–F). The red arrows indicate the six divergent regions identified in Figure 2, the blue arrow in (F) indicates the location of the *CORIN* gene (Figure 5).

Having genomic data from all three taxa in the genus allowed us to estimate the three-taxon PBS values for each of the individual *Vermivora* species. *V. bachmanii* showed very high lineage-specific evolution, with high background PBS values and large (i.e. >>5Mb) regions of higher divergence clustered in chromosomal locations with presumed low recombination rates (e.g. the Z chromosome; Irwin 2018). By contrast, and as expected based on low *F*_ST_ values, PBS values among *V. cyanoptera* and *V. chrysoptera* were very low, with the only exception between the regions housing pigmentation genes.

We compared PBS for *V. bachmanii* to the *F*_ST_ values contrasting *V. cyanoptera* and *V. chrysoptera* for two regions with key pigmentation genes (*ASIP* and *BCO2*), which have been the focus of previous work in *Vermivora* and *Setophaga* Warblers (Toews et al. 2016, Wang et al. 2021, Baiz et al. 2021). Figure 4A shows that *V. bachmanii* has elevated divergence at *ASIP*, at the same region where *V. cyanoptera* and *V. chrysoptera* are divergent. By contrast, for *BCO2* (Figure 4B), there does not appear to be evidence of lineage specific evolution in *V. bachmanii*.

**Figure 4.**
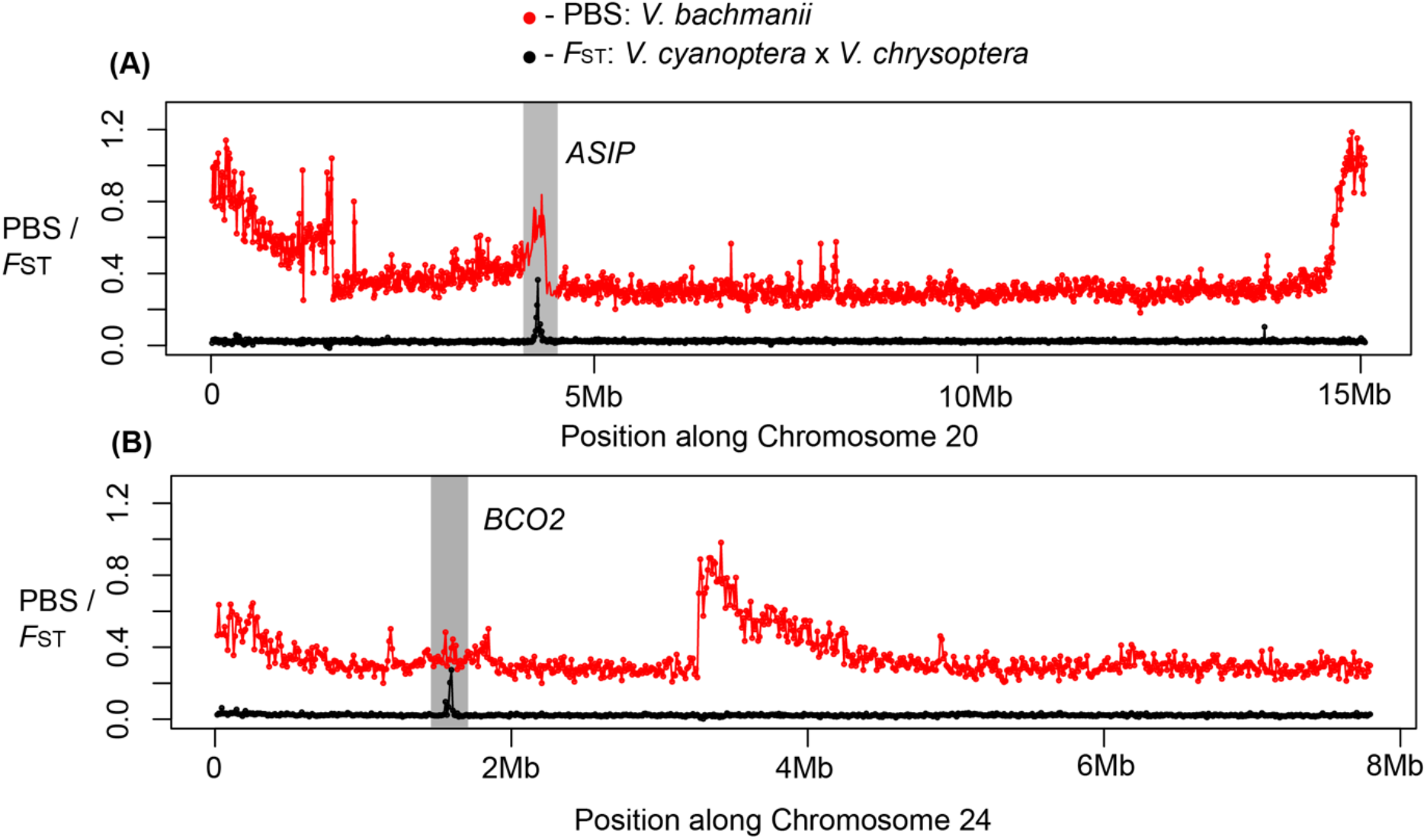
*F*_ST_ between *V. cyanoptera* and *V. chrysoptera* and PBS estimates (in non-overlapping 10kb windows) for *V. bachmanii* for two pigmentation gene candidates (A) *ASIP* and (B) *BCO2*.

One region on chromosome 4 had elevated PBS in *V. chrysoptera* (between 25,545,000 and 25,625,000 bp) but not in *V. cyanoptera* (Figure 5); this region had not been previously reported as being a strong outlier in *F*_ST_ (Toews et al. 2016; Figure 3C), but it was a clear outlier for PBS for *V. chrysoptera* (Figure 3F). This divergent region overlaps only a single gene, *CORIN*, which is a known modifier of *ASIP* and plays a role in coat color in mice (Enshell-Seijffers et al. 2007).

**Figure 5.**
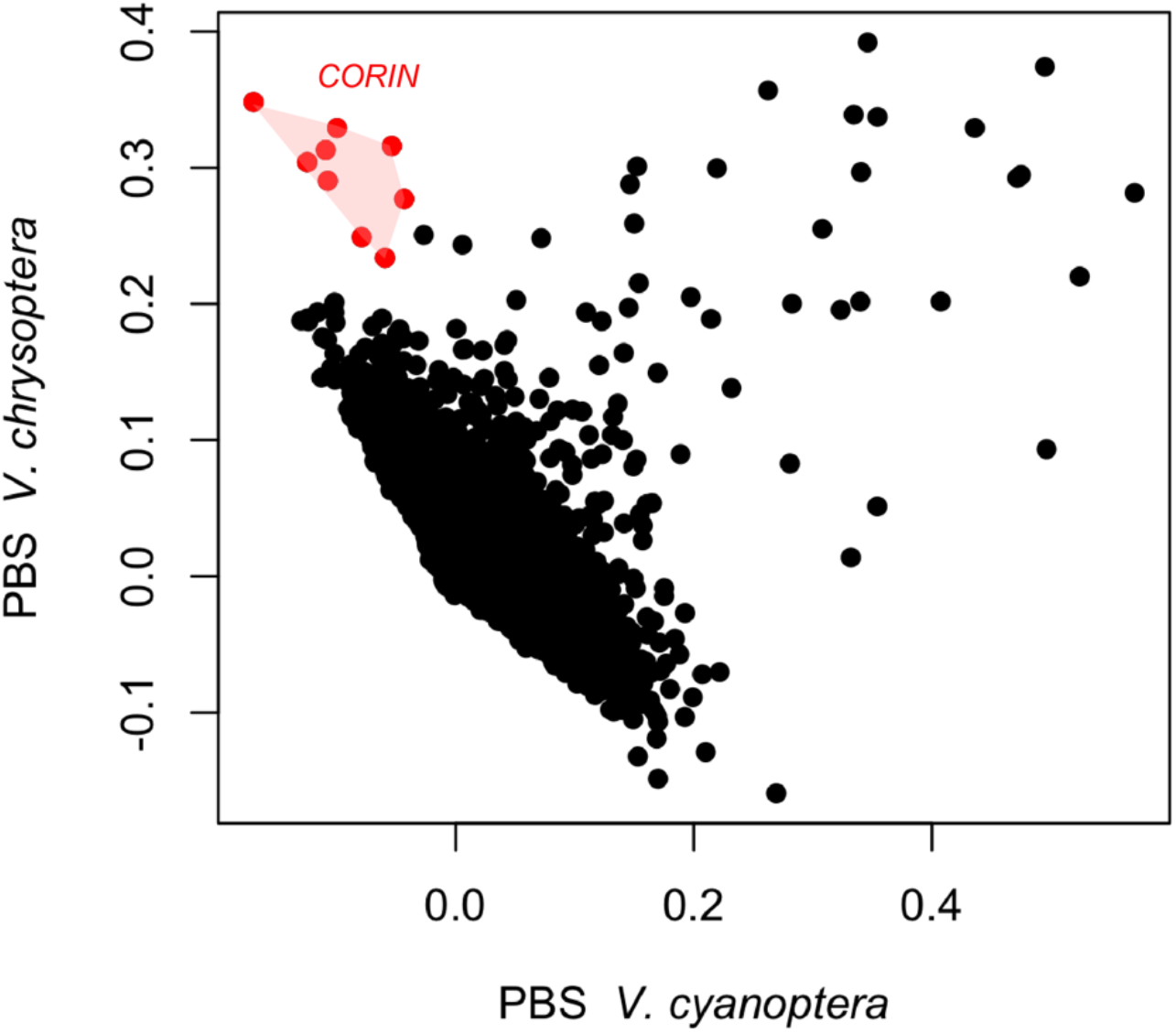
Genome-wide PBS estimates (in non-overlapping 10kb windows) for common windows between *V. cyanoptera* and *V. chrysoptera*. The previously undocumented divergent region in *V. chrysoptera* on chromosome 4 (between 25,545,000 and 25,625,000 bp) with the *CORIN* gene is shown in red.

We restricted our ROH calls to only long ROH (those greater than 1Mbp), as shorter ROH are unlikely to be accurately identified with such sparse data. For each individual we calculate F_ROH_, which is the sum length of all ROH divided by the autosomal genome length (calculated to be 938,591,657 bps), and N_ROH_, which is the total number of ROH segments found in that individual.

We found high levels of total genomic ROH (Figure 6A) in both *V. chrysoptera* and *V. bachmanii*, each significantly higher than *V. cyanoptera* (Wilcoxon rank sum test, two-tailed *p* = 1.638×10^-3^ and *p* = 1.881×10^-2^, respectively). However, we found no significant difference in F_ROH_ levels between *V. chrysoptera* and *V. bachmanii* (Wilcoxon rank sum test, two-tailed *p* = 0.8749), although we do note that the highest F_ROH_ value observed was from *V. bachmanii* (sample AMNH-759216, F_ROH_ = 0.0548).

**Figure 6.**
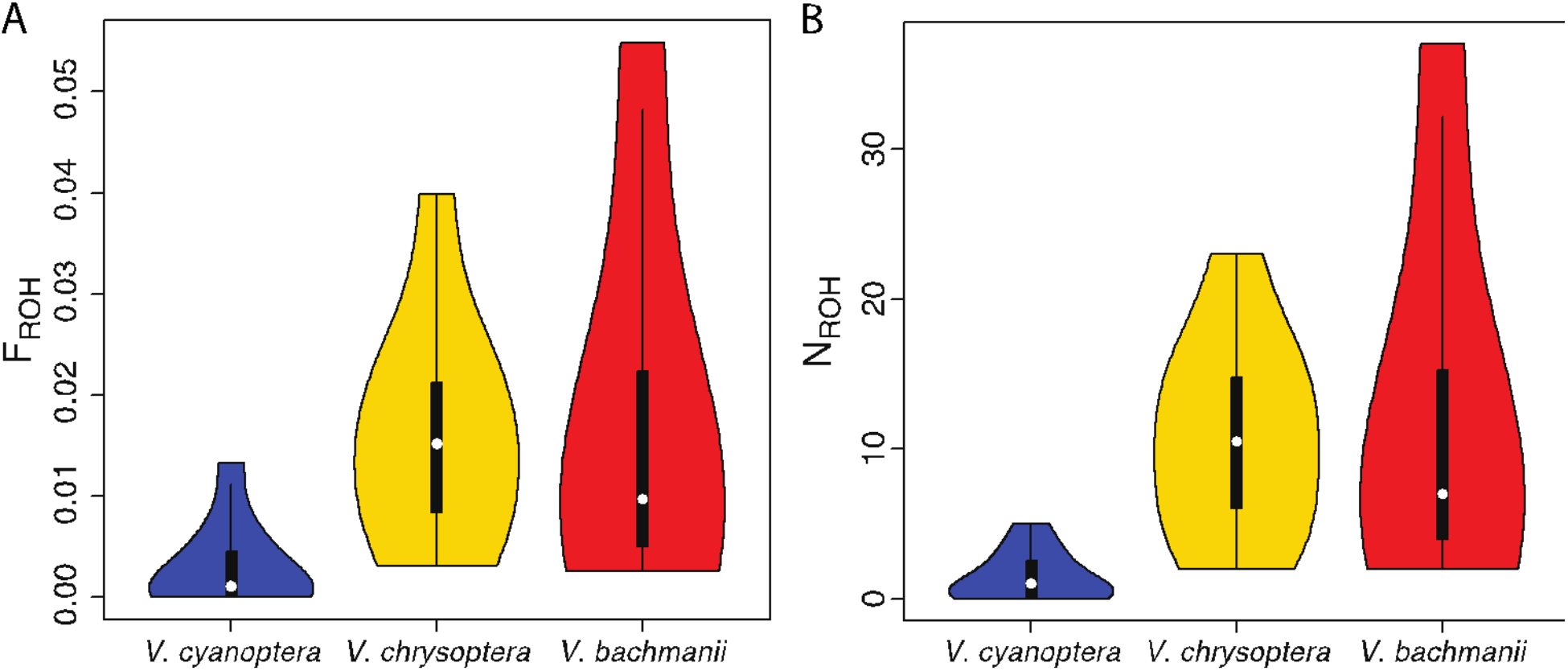
Distribution of (A) F_ROH_, the fraction of the genome covered by ROH, and (B) N_ROH_, the total number of ROH segments, among *V. cyanoptera, V. chrysoptera*, and *V. bachmanii* individuals. Each statistic computed from individual ROH calls ≥ 1Mbp long.

We found similar patterns among the species when comparing the total number of long ROH segments (Figure 6B). Both *V. chrysoptera* and *V. bachmanii* each had significantly higher N_ROH_ than *V. cyanoptera* (Wilcoxon rank sum test, two-tailed *p* = 7.036×10^-4^ and p = 7.162×10^-3^, respectively), but no significant difference between them (Wilcoxon rank sum test, two-tailed *p* = 0.9134). The *V. bachmanii* sample AMNH-759216 again showing highest observed N_ROH_ = 37.

## DISCUSSION

We synthesize these new sequencing data with past knowledge of genotype-phenotype associations in *Vermivora* Warblers. Specifically, we found evidence supporting a long history of divergence and lineage-specific evolution for *V. bachmanii*. We also find possible evidence of inbreeding for both *V. bachmanii* and contemporary *V. chrysoptera*. Finally, these new data also allowed us to quantify novel divergence patterns, and in *V. chrysoptera* one of these divergent regions was associated with a previously undocumented melanogenesis pigmentation gene candidate.

Across the divergent regions between *V. chrysoptera* and *V. cyanoptera, V. bachmanii* consistently fell into its own cluster along the first PC axis, which reliably explained disproportionately more variation (>24%) than all other PC axes. Unlike the contemporary hybrids, which fell between *V. chrysoptera* and *V. cyanoptera* along PC2, *V. bachmanii* showed no evidence of such intermediacy, which would be consistent with evidence of admixture and/or introgression at these gene regions. Moreover, the non-significant ABBA-BABA test across the genome implies no evidence of historical admixture. Thus, while there are phenotypic similarities between *V. bachmanii* and some of the contemporary *Vermivora* hybrid phenotypes (e.g. “Lawrence’s Warbler”), our data suggest this similarity is not likely the result of introgression at previously identified pigmentation genes (e.g. *ASIP*). Indeed, all the genomic evidence is consistent with *V. bachmanii* having been a highly divergent, reproductively isolated species.

That said, it is likely that parallel evolution at common gene targets are responsible for some of the phenotypic similarity across the genus. For instance, the PBS analysis shows that *ASIP* has elevated lineage-specific evolution in *V. bachmanii* (maximum PBS for the ASIP region: *V. bachmanii =* 0.84, *V. chrysoptera* = 0.30, *V. cyanoptera* = 0.15) and this is consistent with selection at this gene for *V. bachmanii’s* distinct melanic traits. This is a common pattern with *ASIP*: multiple avian lineages, including other warbler species showing melanin pigment differences, show clear evidence of repeated, independent divergence at *ASIP* (Campagna et al. 2017, Wang et al. 2020, Baiz et al. 2021, Semenov et al. 2021, Campagna and Toews 2022). We note, however, the distribution of melanin in *V. bachmanii* throat patches also appears qualitatively different from *V. chrysoptera*, with some *V. bachmanii* patches extending fully down the breast to the belly (Fig 1C), implying a different expression pattern.

The patterns of lineage-specific evolution at *BCO2* across the three *Vermivora* speaks to the broader evolution of this gene in this genus. First, unlike *ASIP*, there is no elevated PBS for *V. bachmanii* in the *BCO2* region. For the extant *Vermivora*, the PBS values are approximately twice as high in *V. chrysoptera* than *V. cyanoptera* (maximum *V. chrysoptera* PBS for chromosome 24=0.25, maximum PBS for *V. cyanoptera* = 0.11). The evolution of *BCO2* is thought to be linked to the distribution of body carotenoids, which is extensive for both *V. cyanoptera* and *V. bachmanii*, but not *V. chrysoptera. BCO2* degrades carotenoids, therefore the reduced action of its gene product relates to an increase in observable yellow carotenoids in the feathers (Gazda et al. 2020). Taken together, these PBS results suggest that the broad distribution of yellow coloration was possibly the ancestral state in *Vermivora*, with higher lineage-specific evolution in *V. chrysoptera* for the derived state of reduced carotenoid distribution (i.e. higher *BCO2* expression). RNA expression information from the developing feathers of extant *Vermivora* warblers would be required to test this further.

The inclusion of *V. bachmanii* in the PBS analysis facilitated the discovery of entirely novel pigmentation gene candidates in *Vermivora* (and parulids more generally). Specifically, the elevated lineage-specific evolution on chromosome 4 in *V. chrysoptera* (Figure 5) was not previously identified as a strong outlier in *F*_ST_ with *V. cyanoptera* (Figure 3C). The maximum PBS in this region for *V. chrysoptera* is 0.35, whereas the maximum value is −0.04 in *V. cyanoptera*. This divergent region overlaps with only a single gene, *CORIN*. The gene product of *CORIN* is a modifier of the “agouti pathway” and, in mice, acts downstream of *ASIP* as a suppressor (Enshell-Seijffers et al. 2007). In *Vermivora* warblers, variation in the promoter region of *ASIP* has been linked to melanic variation in throat and mask color in *V. chrysoptera* (Toews et al. 206; Baiz et al. 2021). *CORIN* has also been implicated in the melanogenesis pathway of other birds, like crows (Poelstra et al. 2015) and *Zosterops* white-eyes (Bourgeois et al. 2016), as well as pelage color variation in big cats (Xu et al. 2017). In mice, binding of agouti to *MC1R* reduces its activity and results in a shift from the production of eumelanin (black) to the synthesis of pheomelanin (yellow; Enshell-Seijffers et al. 2007). These studies point to the possible action of *CORIN* as downstream of the agouti signaling pathway and it acts to suppress agouti protein activity. Thus, increased *CORIN* expression in *V. chrysoptera* could result in suppression of *ASIP* and, subsequently, higher melanin expression in the throat and mask. Again, expression data will be critical to collect to test these new hypotheses.

Our analysis of long runs of homozygosity (ROH) suggested that both contemporary *V. chrysoptera* and historical *V. bachmanii* may have experienced a small effective population size or a population bottleneck and possibly inbreeding. The timing of a bottleneck or inbreeding is unclear, but demographic modeling of *V. bachmanii* showed that the species exhibited the expected pattern of population expansion in the Late Pleistocene (Smith et al. 2021). ROH can harbor increased levels of deleterious homozygotes (Szpiech et al. 2013; Szpiech et al. 2019), and therefore their prevalence is important for studies of inbreeding depression in wild (Stoffel et al. 2021a; Stoeffel et al. 2021b; Nguyen et al. 2022) and domesticated (Curik et al. 2014; Mooney et al. 2021) animals, with potential implications for the long-term survival of the population/species. Whether the pattern among extant *Vermivora* indicates that *V. chrysoptera*’s fate may be similar to that of *V. bachmanii* is unclear. At least among extant *Vermivora*, population declines of *V. chrysoptera* have been the most concerning, particularly across the Appalachian populations (Rosenberg et al. 2016; Karmer et al. 2017). However, an important caveat to consider in the current analysis, given the low-sequencing coverage of these samples, is that ROH calling is likely to be quite noisy. Given that these samples were all sequenced to approximately the same coverage and processed with the same filters, we expect that these ROH patterns are at least comparable among the samples analyzed here, but suggest that future, higher-coverage genomic studies should carefully investigate these patterns further.

Taken together, our study and others like it spotlight the invaluable stores of knowledge that rest in natural history collections like the CUMV and AMNH, while also highlighting how the extinction of *V. bachmanii* represents the loss of an evolutionarily divergent, fully reproductively isolated taxon. In 1890, C.S. Gailbraith, the researcher that collected the one of Bachman’s Warbler sequenced here (Figure 1C), likely did not anticipate its pending extinction. Moreover, he could never have imagined the tools available to extract new knowledge from its tissues. Investments in museums return healthy dividends in normal times; in the midst of the Anthropocene extinction the value of such investments are incalculable.

## ACKNOWLEDGEMENTS

We thank both the CUMV and AMNH for access to specimens. We thank Jennifer Walsh for assistance with the toepad DNA extraction at the CUMV and for comments on an earlier version of this manuscript. This work was supported by NSF-DEB (award# 2131469 to DPLT) and NSF-DBI (award# 2029955 to BTS), start-up funds from Huck Institutes of the Life Sciences and the Pennsylvania State University’s Eberly College of Science (DPLT and ZAS). Computations for this research were performed using the Pennsylvania State University’s Institute for Computational Data Sciences’ Roar supercomputer.

